# Identifying the multiple drivers of Cactus diversification

**DOI:** 10.1101/2023.04.24.538150

**Authors:** Jamie B. Thompson, Tania Hernández-Hernández, Georgia Keeling, Nicholas K. Priest

## Abstract

Many drivers of diversification have been identified across the angiosperm Tree of Life, ranging from abiotic factors, such as climate change, to biotic factors such as key adaptations. While this provides invaluable evolutionary insight into the rise of major angiosperm lineages, our understanding of the complexity underlying this remains incomplete. In species-rich families such as Cactaceae, simple explanations of triggers of diversification are insufficient. Their sheer morphological and ecological diversity, and wide distribution across heterogeneous environments, render the identification of key forces difficult. Cactus diversification is likely shaped by multiple drivers, which themselves interact in complex ways. This complexity is extremely difficult to disentangle, but applying modern analytical methods to extensive datasets offers a solution. Here, we investigate the heterogeneous diversification of the iconic Cactus family. We reconstruct a comprehensive phylogeny, build a dataset of 39 abiotic and biotic variables, and predict the variables of central importance to tip-speciation rate variation using Machine Learning. State-dependent diversification models confirm that a rich range of eleven abiotic and biotic variables filtered as important by Machine Learning shape Cactus diversification. Of highest importance is an atypical latitudinal gradient in speciation rates, which is spatially decoupled from richness hotspots. Of medium importance is plant size, shaped by growth form. Of lesser, but significant, importance is soil composition, bioclimate, topography, geographic range size, and chromosome count. However, it is unlikely that any one of these eleven variables is of primary importance without the complex interactions formed with others. Our results contribute to our understanding of one of the most iconic angiosperm families, while revealing the need to account for the complexity underlying macroevolution.

## Introduction

The angiosperm family Cactaceae is an iconic component of ecosystems spanning the Americas (Hernandez-Hernandez et al, 2011; Hernandez-Hernandez et al, 2014). Nearly all Cacti exhibit the succulent life form which enables survival in the face of water scarcity, through adaptations including succulent stems, modified spines and crassulacean acid metabolism (CAM) photosynthesis (Griffiths and Males, 2017). Although Cacti are found across diverse ecosystems including wet tropical forests and colder regions (Pillet et al., 2022), they reach highest richness in arid and semi-arid regions (Gibson and Nobel, 1986; Hernández-Hernández et al., 2014). This has implicated aridification as the central driver of diversification in Cacti, and succulents generally (Gibson and Nobel, 1986; Arakaki et al., 2011). However, aridification cannot explain dramatic within-family phylogenetic imbalances at different taxonomic levels (Hernández-Hernández et al., 2014; Guerrero et al., 2019). For example, subfamilies Maihuenioideae and Pereskioideae contain only ∼19 species in total, whereas over 900 species are found in Cactoideae, while the Mammilloid complex alone holds up to ∼200 species (Breslin et al., 2021). Thus, drivers other than aridification must play a key role in shaping rate heterogeneity within Cacti.

The extraordinary ecomorphological diversity of Cacti implicates biotic mechanisms of diversification, with several identified. Growth forms range from small button-like species in genera including *Epithelantha*, to massive columnars such as *Pachycereus* (Anderson 2001), and pollination syndrome varies from bees to moths or bats (Hernandez-Hernandez et al, 2011; 2014). Diversification rates are fastest in species with larger growth forms and derived pollination syndromes (bat, bird and moth pollination (Schlumpberger, 2012). However, there are a range of other potential drivers of diversification suggested by correlations with biodiversity, including geographic distribution (Majure et al., 2012; Hernández-Hernández et al., 2014; Pillet et al., 2022), elevation (Luebert and Weigend, 2014), global temperature (Silva et al., 2018), chromosome number (Lopes et al., 2021), edaphic properties (Ruedas et al., 2006) and climatic variables recently associated with endemism (Trabuco Amaral et al., 2022). Additionally, we do not know whether Cacti are be influenced by variables known to contribute to broad-scale patterns of diversification in succulents, or angiosperms more generally, including topographic complexity (Verboom et al., 2015) and geographic range size (Hernandez-Hernandez and Wiens, 2020). Finally, we do not understand the level of dependence among the variables shaping diversification rates, a problem beginning to be appreciated in macroevolutionary research.

Previous research has often assessed the impacts of drivers of diversification individually, neglecting the complexity underlying macroevolution. This provides an incomplete understanding, with diversification often shaped by a multitude of drivers, which is exemplified in Cacti by the dependent evolution of growth form and pollination syndrome (Hernandez-Hernandez et al., 2014). Establishing whether elevated diversification rate is truly dependent on both variables, or just one, is difficult. This problem is amplified when considering multiple potential drivers. Recent advances in hidden-states models have provided a more sophisticated framework for confirming individual drivers (Beaulieu and O’Meara, 2016), by allowing unobserved traits to explain rate heterogeneity. However these models are very computationally expensive, cannot integrate dozens of variables into one analysis, are not available for continuous variables such as aridity index, and cannot model non-linear relationships. A complete understanding of the complex process of Cactus diversification requires a thorough investigation with a wide sample of potential drivers, while accounting for interactions.

Recent advances in Machine Learning methods make it possible to efficiently filter for important explanatory variables from dozens of potential variables while accounting for interactions among them (Chen and Guestrin, 2016). This has been applied to reveal key factors underlying the evolution of reef fish diversity (Siqueira et al., 2020), and has the potential to explain the extremely complex evolution of diverse families such as Cactaceae. Here, we apply Machine Learning and phylogenetic methods to help explain the mystery of Cactus diversification. We reconstruct an extensive phylogeny containing ∼69% of described species, and a massive dataset of 39 potential predictors of tip-speciation rates. Many of these have been hypothesised to shape diversification or diversity patterns of Cacti, and several have impacts across succulents and angiosperms generally. Using the tree-based Machine Learning classification method XGBoost (Chen and Guestrin, 2016), we ranked the relative importance of drivers while accounting for interactions among them. Following this filtering, which identified significant predictors, we used phylogenetic state-dependent speciation and extinction (SSE) models to confirm their role and reveal the nature of their relationship with rate heterogeneity. Interestingly, our results indicate that Cactus diversification can be explained by an atypical latitude diversity gradient linked to richness hotspots, a plant-size dependency underpinned by growth form variation, and minor impacts of soil composition, bioclimate, topography, geographic range size and chromosome count. Previous hypotheses of pollinator divergence and aridity are not upheld as primarily important. Their effect is likely to be masked by the drivers we identify as relevant here, after accounting for interactions. Our results reinforce the complexity of biological diversification, as well as identify key factors shaping the rise of Cacti.

## Materials and methods

### Supermatrix assembly and phylogenetic reconstruction

We reconstructed a phylogenetic hypothesis of Cactaceae from a supermatrix built with published genetic sequences. Orthologous loci were identified and clustered with the OneTwoTree pipeline (Drori et al., 2018). The resulting taxonomy was checked manually against CITES (Hunt, 2016). We merged clusters of partial sequences with their full sequences, keeping the longest sequence when a species was present in both partial and full-sequence clusters. We added outgroup sequences from Anacampserotaceae, Portulacaceae and Talinaceae with Mafft -- add (Katoh and Frith, 2012) and visually inspected alignments for quality using SeaView (Gouy et al., 2009). Finally, we trimmed poorly aligned positions with trimAl using the command “gappyout” (Capella-Gutiérrez et al., 2009) before concatenation into a supermatrix with AMAS (Borowiec 2016).

We reconstructed a Maximum Likelihood phylogeny using RAxML v8 (Stamatakis 2014), applying a GTR model of nucleotide substitution to each locus partition, and assessing topological support by allowing bootstrapping to end automatically. In this analysis, we constrained the monophyly of subfamilies and the tribe Echinocereeae to improve the likelihood calculation, after initial ML searches. We time-calibrated the final phylogeny under Penalized Likelihood (PL) using treePL (Smith and O’Meara, 2012), with stem and crown ages given upper and lower bounds from the highest posterior probabilities estimated by Ramírez-Barahona et al., (2020). To identify the most common cross-validation optimal parameters we performed 100 priming steps in treePL. Finally, we performed cross-validation to find the smallest score, and estimated divergence times using the corresponding smooth value. More sophisticated methods of time calibration such as BEAST (Drummond and Rambaut, 2007) were not possible given the size and complexity of this dataset, and we used secondary calibrations due to limited fossil availability (Arakaki et al., 2011).

### Data assembly

We compiled a large dataset of abiotic and biotic variables potentially contributing to variation in Cactus diversification rate. Many have been previously hypothesised to drive Cactus diversification (e.g. growth form, pollination syndrome and aridity), while some play a role in succulent or angiosperm diversification generally (e.g. latitude, elevation, bioclimate, plant size, topographic complexity and geographic range size). For spatial variables, we downloaded georeferenced distribution data from the Global Biodiversity Information Facility (GBIF), retaining only those present in our phylogeny, and cleaned coordinates manually. We plotted distributions genus-by-genus and removed coordinates outside described ranges, from multiple sources including online floras and Anderson (2001). With these curated coordinates, we calculated median latitude per species, and extracted the 19 bioclim variables (Booth et al., 2014), aridity index and potential evapotranspiration (Trabucco and Zomer, 2018), elevation (Amatulli et al., 2018), three relevant measures of soil (sand content, water and texture) (Poggio et al., 2021), six measures of topographic complexity (slope, roughness, standard deviation of slope, standard deviation of elevation, profile curvature and tang curvature) (Amatulli et al., 2018), and biome (Guan et al., 2012). We used median values for each species in all further analyses, except biome which is categorical, for which we used the most common biome for each species. We calculated geographic range sizes as Area of Occupancy (AOO) using the R package red (Cardoso, 2017), with a grid size of 2x2km as required by IUCN. We downloaded chromosome counts from the Chromosome Counts Database, using the median for each species in our analyses (Rice et al., 2015). We built upon size (height or length), growth form and pollination data collected by Hernandez-Hernandez et al. (2014). Most of these data were from Anderson (2001), but a small amount were from publications or descriptions of specimens from online databases. Plant size data was mostly only available as minimum and maximum, and we recorded maximum to avoid issues with observations of incomplete, juvenile or diseased specimens. Following Hernandez-Hernandez et al. (2014), we binarised growth form as globose solitary, globose caespitose or barrel form, versus arborescent, shrubby or columnar, recognising the complexity of assigning growth forms to Cacti (Hernandez-Hernandez et al. (2011). Similarly, following Hernandez-Hernandez et al. (2014) we binarised pollination syndrome as ancestral (mellitophily, or bee-pollination) versus derived syndromes (ornithophily (birds), chiropterophily (bats) and sphingophily (moths)). We also recorded whether a species is epiphytic from Anderson (2001), publications or online databases.

### Diversification analyses

We estimated diversification dynamics after pruning outgroups using Bayesian Analysis of Macroevolutionary Mixtures (BAMM) (Rabosky, 2014), sampling four Metropolis coupled Markov-chains (MCMC) of 50 million generations every 5,000 and discarding the first 10% as burn-in. We set priors with the R package BAMMtools (Rabosky et al., 2014), and implemented a conservative prior of a single rate shift. To account for imbalanced sampling, we provided sampling fractions for every genus according to CITES (Hunt, 2016). We assessed convergence with the R package coda (Plummer et al., 2006), ensuring effective sample sizes were >200. Mean tip speciation rates were estimated with BAMMtools. With mean tip speciation rates estimated by BAMM as the response variable, we assessed the relative importance of variables using the tree-based Machine Learning classification method XGBoost (Chen and Guestrin, 2020). XGBoost assesses the importance of hypotheses explaining diversification by applying an adaptive learning algorithm to a set of models that are progressively better fit to the data by reweighting extreme residuals of the previous model. XGBoost provides benefits over traditional methods for this type of problem, such as phylogenetic generalised least squares which has been used to analyse family-level dynamics in plants (Hernandez-Hernandez and Wiens, 2020). Importantly, these benefits include relaxing the assumption of linear relationships between variables and the response, capturing complex interactions in high-dimensional datasets, and better handling of missing data and outliers. By adapting R code from Siqueira et al. (2020), our XGBoost models were tuned in two steps to identify the parameters that minimise the root mean square error (rmse), for the predictive stage. First, an initial tuning step with predefined parameter combinations was performed. Following this, we refit 1,000 models by randomly sampling parameters from uniform distributions bounded by the identified optimal values from stage one +/- 10%. For the final predictive model, the combination of parameters that minimised rmse was used, and tree-depth varied from 2-7. A cross-validation procedure assessed the accuracy of the XGBoost in predicting tip-rates, by randomly subsetting 80% and 20% of the data into training and testing parts, respectively. We refit the final model using the training dataset and then used the coefficients of prediction to predict tip-rates in the testing dataset. To explicitly test whether complexity better explains Cactus diversification than simple models, we reran this test but did not allow tree-depth to vary from one, and compared mean R^2^. After visually inspecting BAMM-estimates tip-rates and the explanatory variables, we performed a sensitivity test. Although XGBoost is robust to outliers, we wanted to ensure our results were robust and that we uncovered macroevolutionary trends rather than spurious relationships driven by rapidly diversifying lineages. Four genera had fast tip-rates above 0.65 (*Copiapoa*, *Gymnocalycium*, *Harrisia* and *Pilosocereus*), which could be unlinked to any overarching macroevolutionary trend across the family. These rates could lead to recovering spurious relationships with drivers. We re-ran XGBoost allowing tree-depth excluding all taxa for which tip-speciation rate exceeded 0.65, leaving 996 taxa for analysis, to verify that these rapidly speciating lineages had minimal impacts on results.

### State-dependent diversification models

We used SSE models to confirm the impact of drivers identified as significant by XGBoost (Tables 1 and 2, Figures 3 and 5). We used QuaSSE to analyse the impact of continuous variables on speciation rate (FitzJohn, 2010), after transformation to improve normality (square root elevation, log slope, log soil sand content, absolute latitude after visually inspecting the BAMM estimates). In each analysis, we fit seven models of trait-dependent diversification, in which the relationship between diversification and the variable is constant, linear, sigmoidal and hump (all with-and-without drift, except constant). Models were estimated under maximum likelihood in the R package diversitree (FitzJohn, 2012). We used BiSSE and HiSSE to assess the impact of binary traits on diversification rate (Maddison et al., 2007; Beaulieu and O’Meara, 2016). These methods are similar, however, HiSSE considers more robust null hypotheses when assessing binary-state dependent diversification than BiSSE. BiSSE considers a null model of symmetrical diversification rate variation whereas HiSSE allows rate variation to be driven by an unobserved trait. We ran 10,000 BiSSE MCMC generations in diversitree (FitzJohn, 2012), of a model in which all parameters are free, after setting priors from Maximum Likelihood models. We compared the Maximum Likelihood fits of a free model versus a model in which speciation rates were equal. We did not test models of equal extinction, because they were estimated as equal in the full model. In analyses with HiSSE, we compared a model in which diversification rate changes independently of the focal trait, a trait-dependent model without hidden states, and a trait-dependent model with hidden states.

**Table 1:** Model comparisons of QuaSSE analyses of continuous drivers with significant predictive ability in the XGBoost model, with AIC scores of the best-fitting and null models. Delta AIC is >4 for all bets-fitting models versus second best-fitting models with p<0.0001 for each of the best-fitting models.

| **Trait** | **Best-fitting model** | **ΔAIC of best fitting model versus constant model (null)** |
| --- | --- | --- |
| **Absolute latitude** | Modal with drift | 219.16 |
| **Bio2** | Modal with drift | 261.10 |
| **Bio18** | Modal with drift | 238.06 |
| **Elevation** | Sigmoidal with drift | 220.44 |
| **Profile curvature** | Modal | 369.63 |
| **Geographic range size** | Modal with drift | 307.86 |
| **Size (plant height or length)** | Modal with drift | 76.72 |
| **Slope** | Sigmoidal with drift | 255.29 |
| **Soil sand content** | Modal with hump | 62.9 |

**Table 2:** Model comparisons for BiSSE analyses of discrete drivers with significant predictive power in the complex XGBoost model.

| Model | AIC and ChiSq | P value |
| --- | --- | --- |
| Growth form rates different | 6904.1 |  |
| Growth form rates equal | 6913.1,11.04 | <0.001 |
| Chromosome count rates different | 2716.6 |  |
| Chromosome count rates equal | 2859.6, 144.99 | 0 |

## Results

### Heterogeneous diversification of Cactaceae

We assembled a phylogenetic hypothesis for Cactaceae with 1,069 Caryophyllales species, including six outgroups, using Maximum Likelihood (based on a supermatrix of a 16.47kb alignment from 15 plastid and 3 nuclear nucleotide loci representing 6,020 parsimony-informative sites and 77.5% missing data). Our resulting phylogeny is moderately well-supported, with 32.3% of internal nodes supported by >70% BS and 11.9% by >90% BS (Figure 1). The topology and divergence estimates were broadly congruent with previous hypotheses of Cactus evolution (Araraki et al., 2011; Hernández-Hernández et al., 2014). We estimate the stem age of Cactaceae at ∼48.51 Mya, and the divergence of *Leuenbergeria* and the remainder of the family at ∼37.24 Mya (the crown age). The estimated stem ages of each major lineage are similar, with *Pereskia* at 36.99 Mya, Maihuenioideae and Opuntioideae at a near-simultaneous 36.86 Mya and Cactoideae at 35.76 Mya. Estimated crown ages of *Leuenbergeria*, *Pereskia*, Maihuenioideae, Opuntioideae and Cactoideae are 19.65, 34.94, 4.03, 17.13 and 35.76 Mya, respectively.

**Figure 1:**
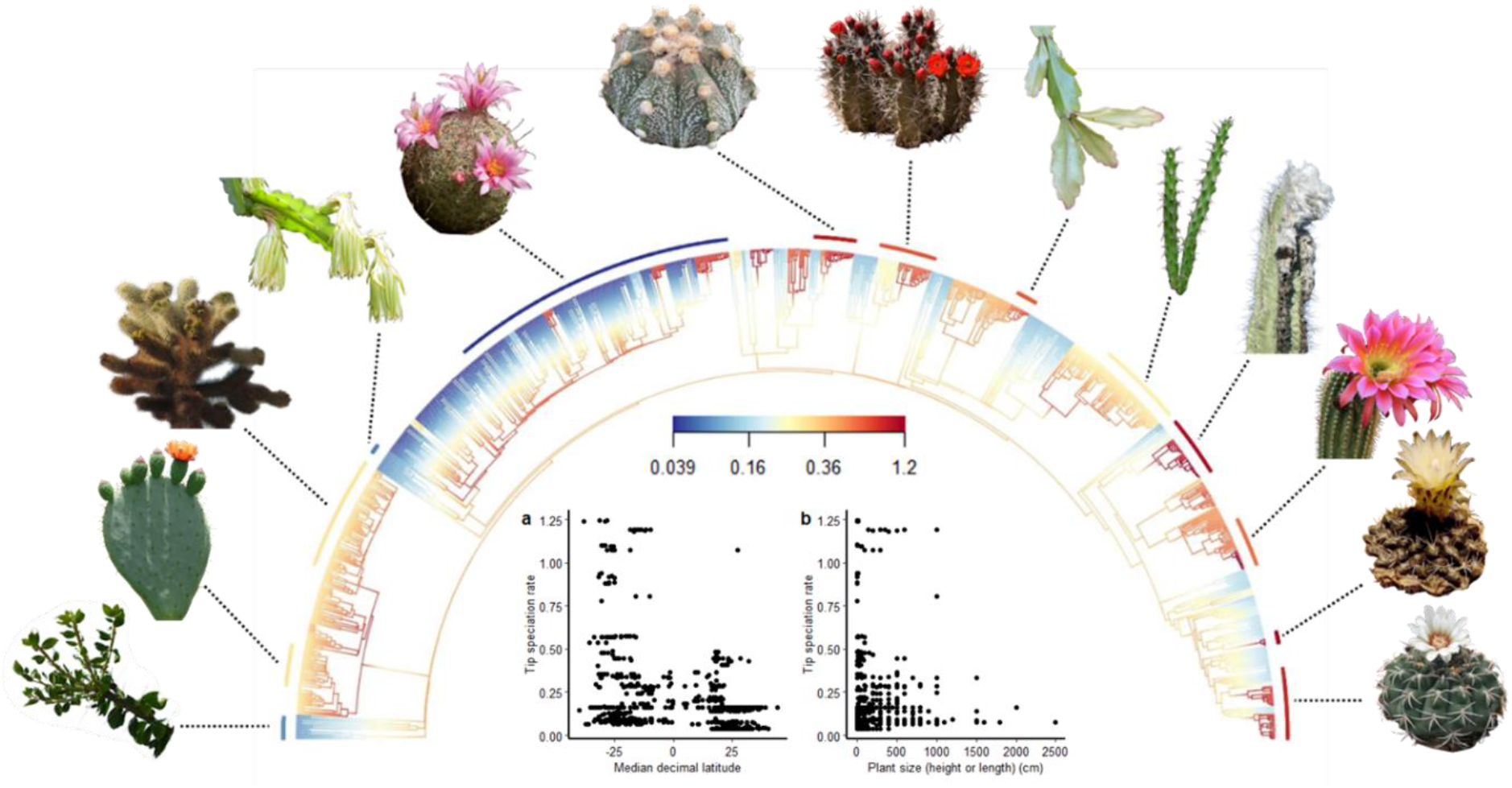
Remarkable diversification rate heterogeneity across Cactaceae is shaped strongly by latitude and size (height or length). BAMM-estimated speciation rates vary 32-fold across the family. Arc segments of median speciation rate for thirteen morphologically varied Cactus genera are indicated (edited images are from Flickr and are available under a Creative Commons licence). Scatterplots show the relationship between tip speciation rates, and median latitude and size (height or length).

Diversification rate heterogeneity is remarkably varied in Cactaceae (Figure 1). We recover 32-fold variation in speciation rate, with strong support for rate heterogeneity (Bayes Factor 4096) and with the best-supported rate shift configuration comprising 19 shifts (Supplementary Figure 1). Tip speciation rates varied 32-fold, from <0.04 in some *Mammillaria* to >1.2 in *Gymnocalycium* and *Pilosocereus*. Despite being a rich subfamily, only one shift was detected in Opuntioideae, in the basal branch. In contrast, multiple shifts were detected across Cactoideae, including several within the *Mammillaria* complex, and shifts in tribes Cereeae, Trichocereeae, Cacteae, Hylocereeae and Browningieae.

### Multiple drivers of Cactus diversification

We assembled a dataset of variables for macroevolutionary analyses. This comprised 39 explanatory variables for up to 1,063 ingroup species, with 22.3% missing data and a mean of 826 entries per variable, ranging from 374 (chromosome count) to 1,063 (growth form). The bulk of the data (34 explanatory variables) were derived from the GBIF coordinates which, after genus-by-genus curation, comprised 33,030 location coordinates for 851 species present in the phylogeny. There is varied coverage in the location data, with an average of 39 coordinates available for each species (ranging from one to 943). The full dataset is available in Supplementary Materials.

We identify 11 variables that are significant predictors of tip-speciation rates. There are seven abiotic and four biotic significant predictors in the XGBoost model which accounts for interactions with tree depth varying up to seven (Figure 2a). The primary abiotic driver is latitude (mean predictive power = 0.21, quantiles = 0.20-0.21), with weaker predictive power found for percentage sand in soil (0.04 (0.04-0.05)), precipitation of the warmest quarter (0.03 (0.02-0.04)), elevation (0.03 (0.02-0.03)), mean diurnal temperature range (0.03 (0.02-0.04)), slope (0.03 (0.02-0.04)) and profile curvature (0.03 (0.02-0.04)). The primary biotic variable is plant size (0.12 (0.11-0.13)), with weaker predictive power found for growth form (0.04 (0.03-0.05)), geographic range size (0.03 (0.02-0.03)), and chromosome count (0.03 (0.02-0.04)). This model, which accounted for interactions through varying tree depths, had very high prediction accuracy (mean bias = 0.032) and moderate precision (mean R^2^ = 0.22).

**Figure 2:**
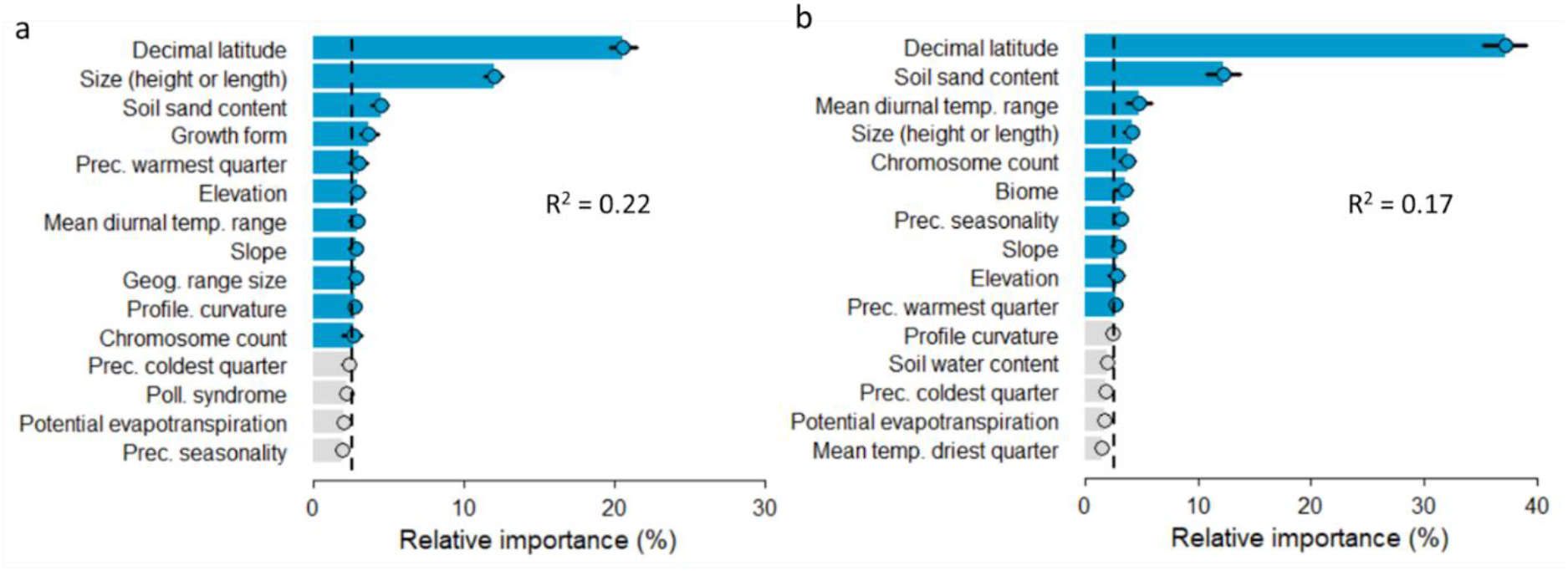
Importance of explanatory variables identified through Machine Learning models. The relative importance of the top 15 (of 39) explanatory variables in predicting speciation rate in 1,000 XGBoost bootstrap replicates is plotted for complex models with maximum tree-depth of seven (a), versus simple models with maximum tree-depth of one (b), with model precision indicated by mean R^2^. The vertical dashed line indicates the threshold of predicting speciation rate by chance expectation alone. When interactions are accounted for, the relative importance of several variables shifts, notably latitude, and the R^2^ increases.

We find that not accounting for complex interactions reduces predictive power and can lead to false inferences in macroevolutionary analysis. Overall, predictive power is lower in the XGBoost model not accounting for interactions (mean R^2^ = 0.17; Figure 2b). Interestingly, the complexity of the model not only alters the rank order of importance on diversification, but can also determine whether individual variables are predicted to influence diversification. Two variables (biome and precipitation seasonality) were identified as significant predictors in the simple model, but are non-significant after accounting for interactions between variables. Other variables were not identified in the simple model, but are significant after accounting for interactions (Figure 2b). Though some of the changes are small, the rise in importance of growth form as a predictor of tip-speciation rates with the inclusion of complex interactions is notable (from 0.0027 (0.0019-0.0034) to 0.04 (0.03-0.05)).

Results of the complex model are relatively robust to outliers. XGBoost accounts for outliers, nevertheless, we replicated analyses without species with tip-speciation rates above 0.65. Strong predictive power is maintained for latitude, plant size and growth form, and variation between models is only found in the variables with the weakest predictive power (Outlier Sensitivity Model; Supplementary Materials).

### Relationships between drivers and diversification

We found that each of the 11 tip-speciation rate predictors shape Cactus diversity in different patterns (Figures 3 and 5). We analysed continuous variables with QuaSSE. It is important to note that QuaSSE is limited to estimating general trends, and not the true relationship between variables and diversification (FitzJohn, 2010). Some estimates of variances in modal models are so small that we do not report them (Figure 3). Though not pertaining to the focus of our study, for completeness, we report estimates of drift, the directional tendency across evolutionary time. Overall, we find that the best-fitting relationship between latitude (absolute) and speciation is modal, in which speciation rates are fastest (∼2.36) at ∼+/-22.5°, with positive drift (∼0.0051). For plant size, an inverted modal model is best supported, with speciation rates slowest (∼0.07) at ∼45.8cm and most rapid (∼1.47) at smaller and larger sizes, and a negative drift (−0.060). For percentage sand in soil, a modal model is best supported, in which speciation rates are highest at ∼47.63%, with negative drift (−0.0049). For precipitation in the warmest quarter, a modal model is supported, in which speciation rates are highest (∼1.12) at ∼20.69mm, with positive drift (0.13). For both elevation and slope, negative linear models are best supported. However, they specify extremely unrealistic parameters, and the next-best-supported models are plotted and discussed throughout this manuscript. These are negative sigmoidal models which qualitatively match the conclusions of the negative linear models. For elevation, a sigmoidal model in which speciation rates are highest (∼0.44) below elevations of ∼753.26 and lowest above this threshold (∼0.070), with positive drift (0.49). For slope, a sigmoidal model in which speciation rates are highest (∼0.58) on slopes shallower than ∼2.94° and lowest above this threshold (∼0.41), with positive drift (0.47). For mean diurnal temperature range, a modal model is best supported, in which speciation rates are most rapid (∼3.63) when the diurnal temperature range is ∼13.15°, and drift is positive (−0.035). For geographic range size, a modal model is best supported, in which speciation rate is most rapid (∼3.87) at a range size of 0.23 AOO, with negative drift (−0.045). For profile curvature, a modal model without drift is best supported, in which speciation rates are most rapid (∼1.68) at a profile curvature of ∼0.00023, a mid-range value. We analysed the impacts of binary traits with BiSSE and HiSSE. Our BiSSE analyses showed that rates-different models are better supported for both growth form (ΔAIC = 9, p < 0.001) and chromosome count (ΔAIC = 143, p < 0.001) (Table 2). Species with arborescent, shrubby or columnar growth forms speciate more rapidly (mean 1.72, HPD (1.14-2.31)) than species with globose solitary, globose caespitose or barrel forms (0.44 (0.29-0.60)). Species with derived chromosome counts speciate more rapidly (0.34 (0.27-0.42)) than those with the ancestral count of n = 11 (0.023 (0.013-0.033)). However, state-dependent diversification is not supported by HiSSE, which prefers models with unobserved traits shaping diversification rates. Estimated AIC values are lower for HiSSE models with unobserved traits than trait-dependent models (growth form: ΔAIC = 180.16; chromosome count ΔAIC = 76.78), and null models (growth form: ΔAIC = 67.27; chromosome count ΔAIC = 185.51).

**Figure 3:**
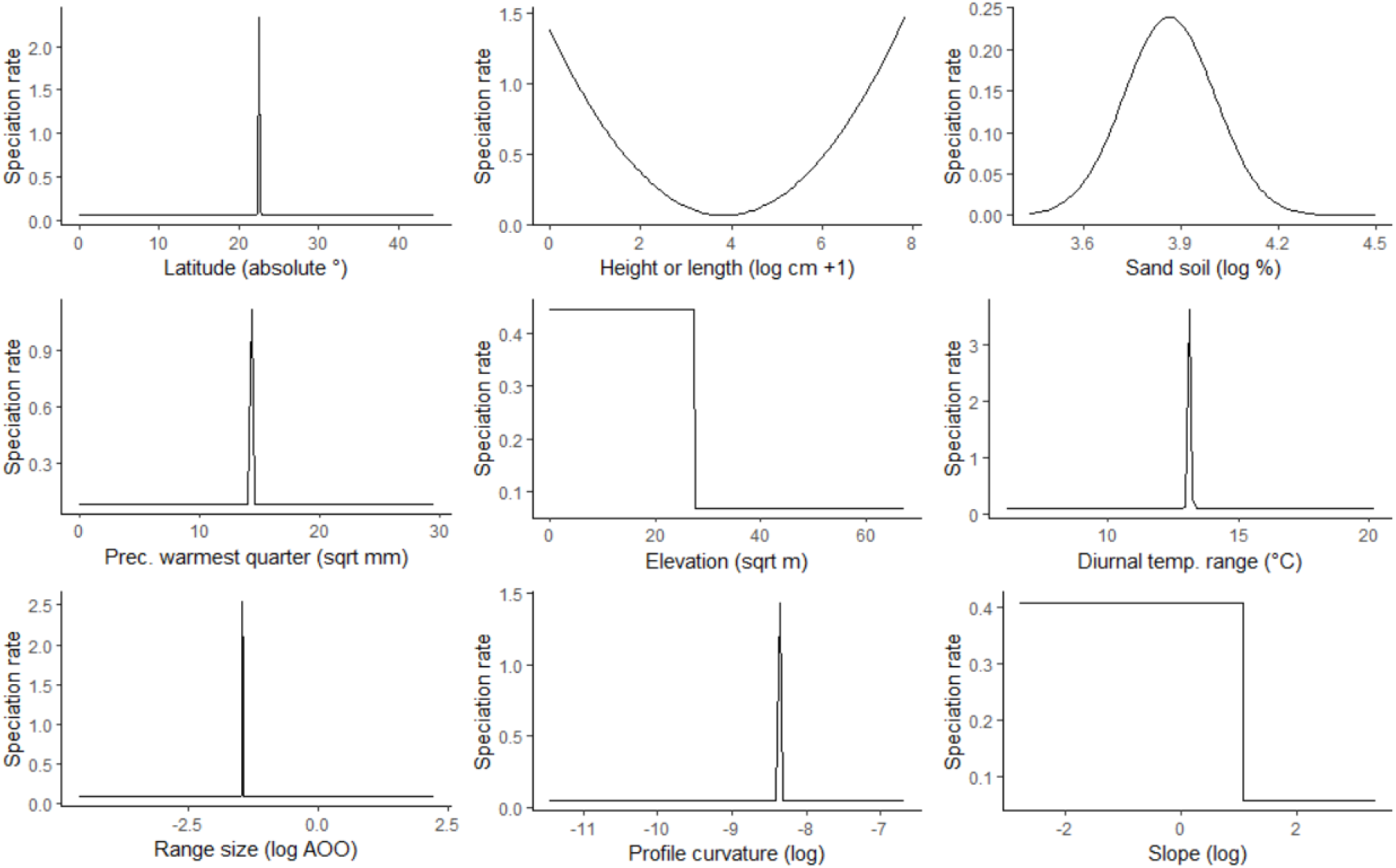
Best-fitting relationships between continuous variables inferred as significant by XGBoost and speciation rate, as estimated by QuaSSE. Variables are in order of ranked importance according to the full XGBoost model. It is important to consider that QuaSSE does not provide the exact relationship between variables and speciation rates, only the general trend (FitzJohn, 2010).

## Discussion

Multiple competing hypotheses have been proposed to explain the diversification of Cactaceae, but it is difficult to extricate key drivers. With a novel phylogeny and extensive dataset of abiotic and biotic variables, we characterise the complexity of Cactus evolution by filtering variables for importance using Machine Learning and confirming the impact of those variables with SSE models. Our analyses support the role of 11 forces shaping diversification in Cacti, including several previously under-investigated drivers, such as geographic range size. We also confer further support for previously hypothesised forces including growth form, although we reveal the impact of unobserved drivers in conferring this effect. Our results reinforce the complexity of macroevolution, with multiple forces acting non-independently to shape diversification, and highlight the necessity for more thorough investigations into lineages across the Tree of Life. These results could improve predictions of how Cactus diversity may fare in the future, an area of active research (Pillet et al., 2022).

### Complexity better explains diversification than simple models

Research into biological diversification across the Tree of Life has long appreciated that macroevolution is extremely complex (Lieberman, 2012; Siqueiro et al., 2020, Givnish et al., 2014, 2015, Vijayakumar et al., 2016, Hernández-Hernández and Wiens, 2020). Though complexity in Cactus diversification has been previously explored by associating radiations with coevolving traits (Hernández-Hernández et al., 2014), the full complexity of factors driving diversification is unknown. Through the implementation of XGBoost, we filtered for the significant predictors of Cactus tip-speciation rates while accounting for interactions amongst predictors. The higher predictive power found in the complex model highlights the importance of extricating such interactions. Failing to account for interactions in the simple model results in the incorrect inference of the relative importance of drivers, such as the reduction in the importance of plant size and the elevated importance of biome. This simple comparison reveals the short-sightedness of failing to consider multiple variables or account for their interactions in shaping diversification rates. Furthermore, the results of hidden-states models reveal that investigating single drivers cannot sufficiently explain diversification. Our results contribute to recent research that has begun to explore complexity in diversification (Beaulieu and O’Meara, 2016; Siqueira et al., 2020) and calls for a more thorough examination across the Tree of Life.

### An atypical latitudinal gradient

QuaSSE predicts peaks in diversification approximately at the Tropics lines of both the Northern and Southern hemispheres. Cacti radiated as they spread from their ancestral Andean range in Chile and Argentina (Hernández-Hernández et al., 2014), and richness hotspots are now found in the American Southwest, Mexico, eastern Brazil and northern Argentina (Pillet et al., 2020). This created an atypical latitudinal diversity gradient (LDG) in which richness peaks at intermediate latitudes either side of the equator, a pattern documented in grasses and attributed to specialisation to cold and arid environments (Visser et al., 2014). Explanations for hotspot generation often present competing hypotheses for the relationship between time in a region and speciation rates (Rahbek et al., 2019). While the cradle hypothesis suggests richness emerges through rapid speciation, the museum hypothesis suggests slow speciation over longer timespans, as seen in the Cape Flora of southern Africa (Verboom et al., 2009).

We find major incongruences between richness and speciation rate in Cacti. We identify hotspots of richness that are consistent with previous research (Pillet et al., 2020), but they exhibit little overlap with estimated hotspots in speciation rate (Figure 4). Though our QuaSSE model predicts peaks in diversification approximately at the Tropics lines of both the Northern and Southern hemispheres (Figure 3), our speciation rate map shows that this Northern Hemisphere pattern is driven by high Cactus species rate in the Caribbean (which is home to *Pilosocereus*, one of the most rapidly-speciating genera in our analysis, see Lavor et al., 2018). Furthermore, in the Southern hemisphere, we identify geographically restricted pockets of speciation, incongruous with regions of high species richness which occur in Eastern Brazil, and Northern Argentina. The otherwise low speciation rate in North America is especially surprising. Mexico is an epicentre of Cactus richness, but our findings indicate that Mexico adheres to the museum hypothesis. Broadly, our results agree with several recent large-scale studies of angiosperms, mammals and marine fishes (Rabosky et al., 2018; Igea and Tanentzap, 2020; Morales-Barbero et al., 2020), but may be driven by a few selected radiations instead of a general trend. While this offers insight into how Cactus richness arose in hotspots, it does not reveal why the subtropical boundaries are optimal for speciation in Cacti. Our finding of hot spots in speciation rate distributed across the Americas highlights why analyses of climatic covariates are needed to resolve the origins of species.

**Figure 4:**
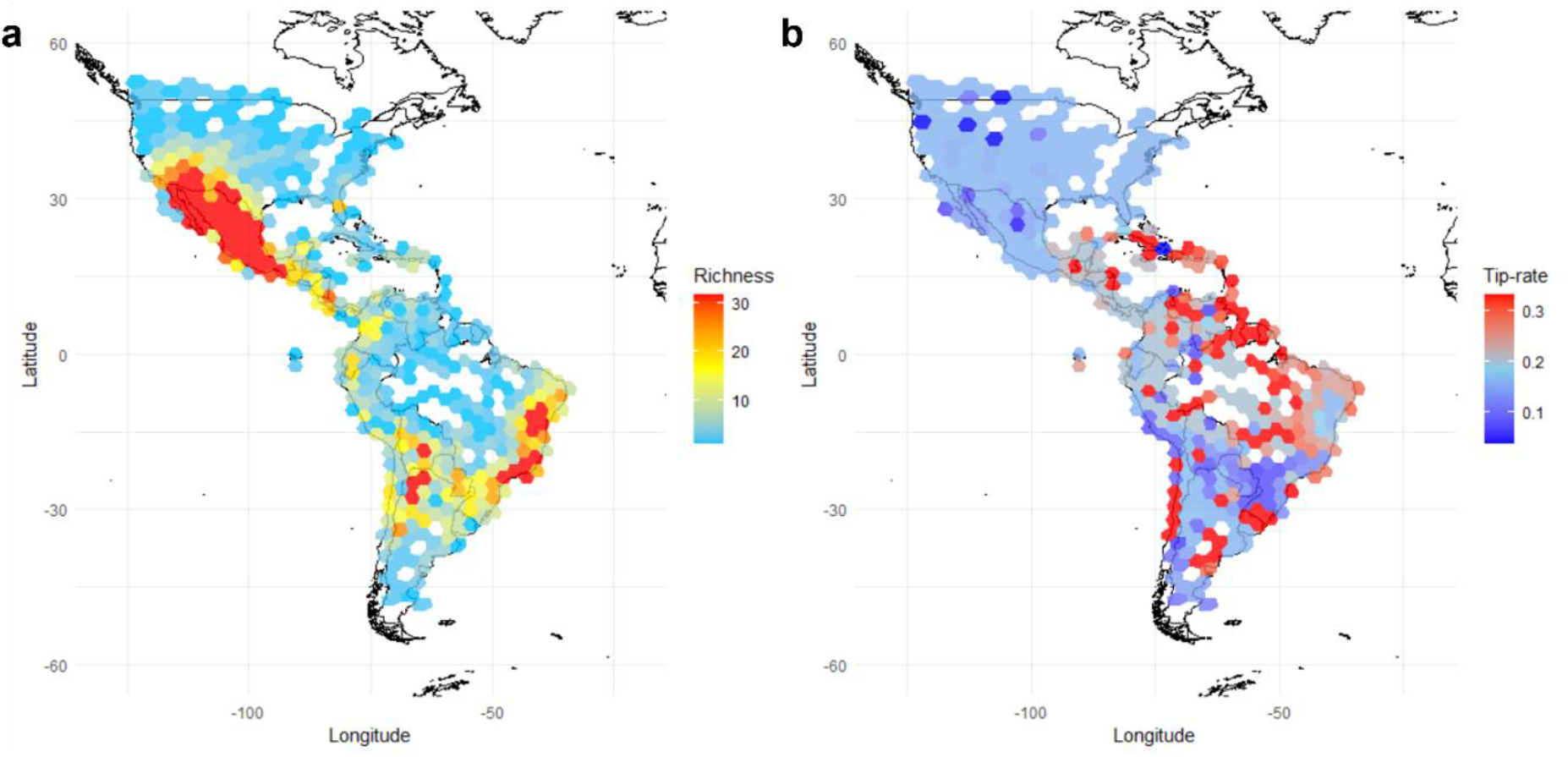
Spatial variation in species richness and speciation rates of Cacti across the Americas. Species richness (a) and median tip speciation rate (b) are estimated for equal-area grid cells (∼322km^2^). Areas of highest richness (e.g. Mexico, Northern Argentina) do not necessarily have the fastest median speciation rates.

### Climatic covariates

Typically, researchers explain LDG patterns with spatial climate-related mechanisms (Mittelbach et al., 2007; Condamine et al., 2012; Rabosky et al., 2018; Brodie and Mannion, 2023; Fenton et al., 2023). However, we find latitude is the strongest predictor in models with-and-without accounting for interactions (Figure 2), indicating the primary role of latitude over covariates. It is important to note that the relative importance of several climatic covariates is reduced in the complex model once interactions are accounted for (Figure 2a). Rapid speciation at both Tropics Lines may simply be explained by geographic expansion promoting rapid diversification in unoccupied ecosystems (Hernández-Hernández et al., 2014). Nevertheless, the Tropics lines remain important because they delimit the centre of the subtropical dry zones (Seidel et al., 2007; Belda et al. 2014). This suggests a role for temporally-varying climatic drivers, which have been linked to Cactus ecology and evolution. Examples include an association between isothermality, temperature seasonality and speciation in *Epithelantha* (Aquino et al., 2021), between diurnal temperature range and CAM photosynthesis (Clifford Gerwick and Williams, 1978), CO_2_ uptake through the seasons (Pimienta-Barrios et al., 1999), and phenotypic variation in species of *Cereus* inhabiting the eastern Brazil hotspot (Trabuco Amaral et al., 2021). Our results support a link between speciation rates and precipitation of the warmest quarter (the growing season) and mean diurnal temperature range.

### Precipitation of the warmest quarter

Precipitation confers selective pressures onto Cacti, and succulent plants generally. The succulent syndrome evolved in order to maximise water storage in regions with relatively low rainfall patterns (Arakaki et al., 2011; Griffiths and Males, 2017), with other adaptations to precipitation including phenology (Cruz and Pavón, 2013). Links between precipitation and diversification have been hypothesised in Cactus previously, in *Epithelantha* (Aquino et al., 2021) and *Eulychnia* (Merklinger et al., 2021), and across the Tree of Life generally (Antonelli, 2017). Precipitation of the warmest quarter, the growing season, predicts over 90% of species richness distribution patterns of the Southern African savannah (Gwitira et al., 2013). Richness begins to increase rapidly at ∼200-250mm, spanning the optimal precipitation for Cactus diversification (∼206.95mm) but plateaus with elevated precipitation. The mechanism by which this level of precipitation in the growing season drives reproductive isolation in Cacti is unclear, but constraints at either extreme may be informative, especially related to growth rates. During the cold season, photosynthetic activity decreases (Sohoulande Djebou et al., 2015). But during the growing season, precipitation and increased temperatures induce germination and growth (Ernest et al., 2000). Many Cacti are very slow growing, which can be especially slow in arid regions (Hultine et al., 2018), reducing generation times. Although a link between growth rate and precipitation in July has been found in *Carnegiea* (Drezner, 2005), the extent to which the predictive power of seasonal precipitation on growth rate extends across Cactus genera is not clear (Delgado-Fernández et al., 2016). Higher rainfall during the growing season may confer increased survivability for non-succulent plants, causing increased competition and resulting in reduced Cactus diversification. Regardless of the mechanisms underpinning this process, it is evident that precipitation of the warmest quarter shapes Cactus diversification. This is broadly consistent with existing research, that has shown precipitation of the warmest quarter predicts over 90% of species richness distribution patterns of the Southern African savannah (Gwitira et al., 2013). Additionally, we find further agreement with the central hypothesis that the succulent syndrome evolved in order to maximise water storage in regions with relatively low rainfall patterns (Arakaki et al., 2011; Cruz and Pavón, 2013; Griffiths and Males, 2017).

### Mean diurnal range

Adaptations to temperature variation may explain our finding of a modal relationship between Cactus diversification and diurnal temperature range. Cacti, which reach their highest diversity in arid and semi-arid regions, have adapted to withstand diurnal temperature ranges by minimising water loss. Such adaptations include species of *Mammillaria* regulating impacts of diurnal temperature variation through modified leaves, and *Carnegeia* through thicker stems (Nobel, 1978). Yet, temperature extremes pose distributional limits in Cacti (Godínez-Álvarez et al., 2003). Many Cacti struggle to withstand high temperatures (Gurvich et al., 2017; Nuzhyna et al., 2018; Andrade and Nobel, 1997), with detrimental impacts conferred by heat on photosynthetic performance with 2**°**C of heating (Aragón-Gastélum et al., 2014). Increased temperatures are associated with germination (Seal et al., 2017) and population or range sizes (Benavides et al., 2021; Pillet et al., 2022). Similarly, most succulents struggle to avoid freezing damage at low temperatures (Griffiths and Males, 2017), which strongly defines the distributions of some species. Constraints on reproductive isolation may be relaxed for Cacti adapted to withstand diurnal temperature fluctuations, due to low population densities of competitors from other plant lineages. But upper and lower limits of temperature changes may be detrimental to Cactus populations.

### Topographic variables

#### Elevation

Spatial abiotic variables such as landscape can also have strong impacts on biodiversity. Landscape is one of the most important determinants of spatial diversity patterns in plants at both large and small scales (Kreft and Jetz, 2007; Wang et al., 2009; Moeslund et al., 2013; Valente et al., 2014; Igea and Tanentzap, 2021). We find that elevation, slope and profile curvature were significant predictors of Cactus speciation rate. Elevational richness patterns often follow a unimodal model, with highest richness at lower-middle elevations (Rahbek, 1995), but not all lineages follow this pattern (Guo et al., 2013). Cacti are found at elevations from sea level to the high Andes, up to ∼4,650m (Hoxey and Lowry, 2021), and a study of Bolivian Cacti revealed a diversity peak at ∼1,000m (Kessler, 2000). We find that Cactus speciation rate sigmoidally decreases with elevation (Figure 3). This contrasts with other plant groups, where radiation is associated with mountain uplift (Hughes and Eastwood, 2006; Givnish et al., 2014, 2015; Xing and Ree, 2017), attributed to encountering new niches as well as by fragmenting populations (as in mammals and birds, see Igea and Tanentzap, 2021). Rapid speciation in lowlands has been documented in woodcreepers (Weir and Price, 2011). Though we do not know why speciation rate is slower, it is interesting that higher elevation is generally associated with a slower tempo of evolution, which is also observed in rates of molecular evolution (Bleiweiss, 1998) and life-history variation (Bears and White, 2009).

#### Slope

Though we know that topographic complexity can shape diversity patterns in plants and that topographic slope influences Cactus richness (Diniz et al., 2021), the impact of slope on diversification is underexplored. In angiosperms and vertebrates, slope can shape biodiversity by creating microclimates of temperature and precipitation (Méndez-Toribio et al., 2016), and influencing hydrological flow, accumulation, soil mixture and erosion (Suggitt et al., 2010; Bogaart and Troch, 2006; Amatulli et al., 2018). We find that slope significantly predicts Cactus speciation rate, with a negative sigmoidal relationship (Figure 3). Shallower slopes may promote survival by providing more favourable niches compared to steep slopes, in which soil erosion and water flow will be higher. But the mechanism by which shallow slopes lead to reproductive isolation remains unclear, other than by a hypothesised greater survival rate. Related to slope, profile curvature is a measure of topographic complexity and speciation in Cacti is fastest in relatively non-curving landscapes. Like slope, it is unclear how this may promote reproductive isolation, but relationships between profile curvature and biodiversity have been found in chaparral shrubs (Moody and Meentemeyer, 2001), marine invertebrates (Demopolous et al., 2018), carabid beetles (Yang et al., 2017) and grasshoppers (Li et al., 2012).

#### Sand

Soil properties have an important role in shaping diversity across plants (Moro et al., 2015; Hulshof and Spasojevic, 2020), including in Cacti (Trabuco Amaral et al., 2022), and are known to shape diversification rates in plants (Buira et al., 2020). Our study indicates a strong impact of soil composition on speciation in Cacti. Edaphic variation determines which species can grow and survive in particular regions, and different species have different requirements (Rajakaruna, 2017). Particular conditions may promote survival, reproduction and population growth in well-adapted species, which leads to reproductive isolation through ecological specialisation, with non-optimal conditions being detrimental to macroevolutionary success (Rajakaruna, 2004). Edaphic variables including sand content contribute to diversity patterns in Cacti including in *Neobuxbaumia* (Ruedas et al., 2006), and species inhabiting the Caatinga (Ribeiro-Silva et al., 2016). Sand content is a relevant property for Cacti, associated with poor water retention (Pachepsky and Rawls, 2001) and transport properties (Passioura, 1988), which provides selective pressure for shallow roots in many Cacti. Soil sand content is the third best predictor of Cactus speciation rates (Figure 3), with a modal model estimating optimal speciation at 47.63% and wide variance. Soils with high levels of sand may be too poor at retaining water even for arid-adapted Cacti, which reduces the opportunity for population growth and reproductive isolation. Furthermore, soils with high sand content are likely to be deserts, which may also have extreme precipitation and temperature patterns, which are non-optimal for Cacti (Pillet et al., 2022). Where soils are less sandy, plants without adaptations to sandy soil are more likely to thrive, providing competition and reducing Cactus diversification rate.

### Biotic variables

#### Plant size and growth form

Plant size is the second most powerful predictor of diversification (Figure 2), but this effect appears to be shaped primarily by growth form variation (Supplementary Figure 2). Plant size has many physiological and ecological consequences (Westoby, 1998), showing allometric scaling with numerous life history traits (Boucher et al, 2017), including those with powerful impacts on speciation rates across angiosperms as a whole (Igea et al., 2017). An inverted modal model is best supported, in which speciation rates are most rapid at smaller and larger sizes, with reduced diversification at intermediate sizes (Figure 3). Continuous size in Cactaceae is primarily shaped by growth form (Supplementary Figure 1), which is a weaker significant predictor of diversification rate. QuaSSE cannot fit models with different rates either side of the midpoint, so we limit our discussion to the effects of growth form analysed with BiSSE, which provides more resolution.

Relationships between organismal size and speciation rate are frequently studied across the Tree of Life (Feldman et al., 2015; Boucher et al., 2020; Cooney et al., 2021), but the nature of trends varies. Typically, smaller species are thought to speciate more quickly due to faster mutation rates and generation times, reduced gene flow, and higher selection on more niche axes, leading to reproductive isolation (Boucher et al., 2017). But in Cacti, diversification rates of larger growth forms are elevated (Figure 5). This pattern has been documented previously in subfamily Cactoideae and is explained through pollinator divergences, which are correlated with transitions to larger growth forms (Hernández-Hernández et al., 2014). Pollinator divergence can compensate for reproductive difficulties conferred by arid biomes, in which mate-finding Allee effects are amplified due to low population densities (Gascoigne et al., 2009). Additionally, pollinator divergence can provide barriers to gene flow (Xu et al., 2012), facilitating reproductive isolation (Smith et al., 2007). Bat and bird pollinators deposit a larger amount of pollen. and disperse over longer distances, than ants or bees (the ancestral state) (Fleming et al., 2009). Pollinator syndrome was not recovered as significant by XGBoost, which may be explained by evolutionary dependence on growth form (Hernández-Hernández et al., 2014), and this interaction was controlled for.

**Figure 5:**
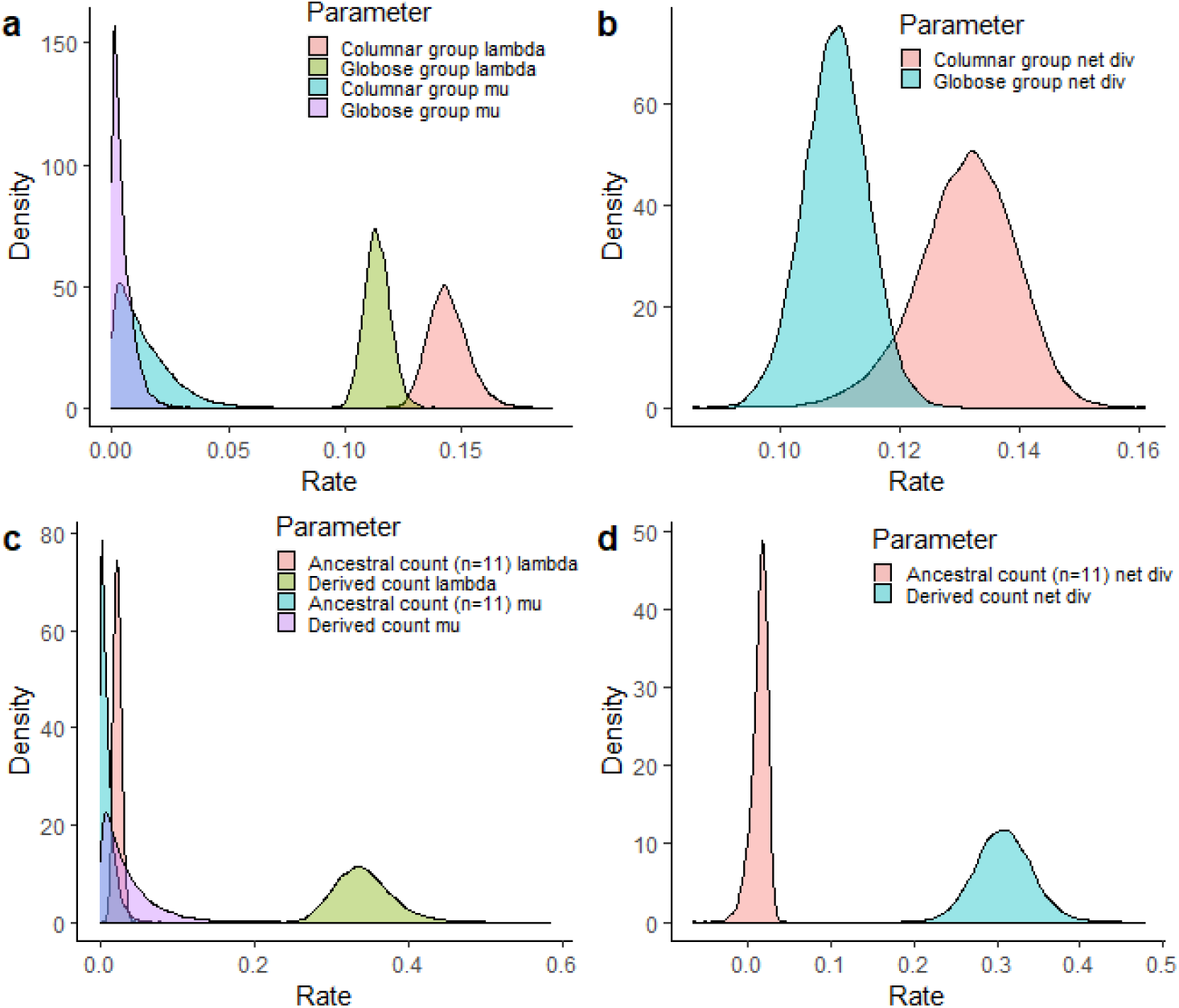
Posterior distribution of speciation and extinction (a and c) and net-diversification (b and d) rates, differing by growth form (a and b) and chromosome count (c and d). Higher rates of species formation are found for Cactus lineages with columnar, shrubby and arboreal growth forms, and lineages with derived chromosome counts. It is important to consider that true state-dependence is not supported in HiSSE models in each trait, and these are visualisations of associations instead of causal relationships.

#### Range size

Geographic range size is a complex biotic variable influenced by a myriad of ecological and evolutionary factors (Sheth et al., 2020). Though range size determines the richness of lineages at the family level in angiosperms, with widespread families diversifying more quickly (Hernández-Hernández and Wiens, 2020), the impact at lower taxonomic levels is poorly understood. We find range size has a weak but significant impact on Cactus speciation, and species with a mid-sized range size speciate most rapidly (Figure 3). Cacti have smaller range sizes than most angiosperms (Goettsch et al., 2015), which are likely to further reduce under climate change projections (Pillet et al., 2022). Smaller ranges confer less resilience to environmental changes and habitat loss (Gaston and Fuller, 2009; Leão et al., 2014), and larger ranges are thought to elevate speciation rate via population fragmentation as the opportunity to encounter barriers and new habitats is magnified (Birand et al., 2012). This pattern is supported by mammals (Cardillo and Bromham, 2003) and birds (Hay et al., 2022). It is hard to reconcile why this pattern was not recovered in Cacti. However, a similar pattern of faster speciation rates of lineages with smaller range sizes shapes the Brazilian flora, attributed to budding speciation (Leão et al., 2020). It is plausible that analysing geographic range size at species level in a rapidly diversifying and young lineage cannot reveal causality. Young species in radiating lineages will necessarily occupy smaller ranges than their ancestors, thus it is hard to estimate the range size at which speciation is most rapid. Nevertheless, understanding the relationship between range size and speciation should be a focus of Cactus research, given the threat of climate change on range sizes (Pillet et al., 2022).

#### Polyploidy

Variation in ploidy has been linked to diversity patterns for decades (Winge, 1917; Stebbins, 1950; Grant, 1981; Rice et al., 2019), with many studies exploring its influence on evolutionary and ecological processes (Ramsey & Schemske, 2002; Sessa, 2019; Zenil-Ferguson et al., 2019). Although the impact on diversification remains highly debated (Kellogg, 2016; Clarke and Donoghue, 2018), several studies have linked polyploidy to radiations across plants (Soltis et al., 2009; Oyston et al., 2022). Chromosome count was the weakest significant predictor of Cactus speciation rate, and derived counts speciate much more rapidly than species possessing the ancestral karyotype (Figures 2 and 5). Cacti have a base chromosome number of 2n = 22, and variation in chromosome number is via polyploidy (Castro et al., 2020). Polyploidy has previously been linked to reproductive isolation in Cactus genera including *Consolea* (Negrón-Ortiz, 2007), *Rhipsalis* (Cota-Sánchez and Bomfim-Patrício, 2010) and *Echinocereus* (Cota and Philbrick, 1994), but no study has investigated this across Cactaceae. Genome duplication can increase diversification rates by creating copies of genes free of selective constraint (Oyston et al., 2022), which can functionally diverge, possibly in response to environmental or ecological change (Van de Peer et al., 2017). Additionally, polyploidy is linked with asexual reproduction and self-fertility (Stebbins, 1950; Manning and Dickson, 1986). Reproductive isolation via polyploidy may promote speciation across Cacti but is unlikely to be a powerful driver of radiation.

## Conclusions

The Cactus radiation remains one of the most iconic lineages of plants (Magallón et al., 2018), especially in arid and semi-arid regions (Arakaki et al., 2011). Cacti are the subject of intense macroevolutionary research, which has revealed a multitude of factors shaping diversification rate, some of which are correlated (Hernández-Hernández et al., 2014). This has made it challenging to identify the important drivers of radiation, a problem found across the Tree of Life (Bouchenak-Khelladi et al., 2015; Donoghue and Sanderson, 2015; Beaulieu and O’Meara, 2016; Hernández-Hernández and Wiens, 2020; Siqueira et al., 2020). Here, we apply a Machine Learning method to efficiently rank the importance of variables in an extensive dataset, revealing eleven key drivers. Cactus speciation is primarily shaped by latitudinal distribution, with regions of rapid speciation broadly decoupled from richness hotspots, a pattern previously demonstrated in many other clades. Morphological variation is also highly important, with larger forms speciating rapidly, contrary to much research on size evolution across the Tree of Life. Further minor contributions are made by soil conditions, topography, geographic range size, bioclimatic variables, and polyploidy.

